# Using Single Nucleotide Variations in Single-Cell RNA-Seq to Identify Subpopulations and Genotype-phenotype Linkage

**DOI:** 10.1101/095810

**Authors:** Olivier Poirion, Xun Zhu, Travers Ching, Lana X. Garmire

**Affiliations:** Epidemiology Program, University of Hawaii Cancer Center, Honolulu, HI 96813, USA; Molecular Biosciences and Bioengineering Graduate Program, University of Hawaii at Manoa, Honolulu, HI 96822, USA

**Keywords:** Single cell, RNA-Seq, single nucleotide variation, cancer, heterogeneity, subpopulations, genotype, linear model, visualization

## Abstract

Despite its popularity, characterization of subpopulations with transcript abundance is subject to a significant amount of noise. We propose to use effective and expressed nucleotide variations (eeSNVs) from scRNA-seq as alternative features for tumor subpopulation identification. We developed a linear modeling framework, SSrGE, to link eeSNVs associated with gene expression. In all the datasets tested, eeSNVs achieve better accuracies than gene expression for identifying subpopulations. Previously validated cancer-relevant genes are also highly ranked, confirming the significance of the method. Moreover, SSrGE is capable of analyzing coupled DNA-seq and RNA-seq data from the same single cells, demonstrating its value in integrating multi-omics single cell techniques. In summary, SNV features from scRNA-seq data have merits for both subpopulation identification and linkage of genotype-phenotype relationship. The method SSrGE is available at https://github.com/lanagarmire/SSrGE.

## Introduction

Characterization of phenotypic diversity is a key challenge in the emerging field of single-cell RNA-sequencing (scRNA-seq). In scRNA-seq data, patterns of gene expression (GE) are conventionally used as features to explore the heterogeneity among single cells^1–3^. However, GE features are subject to a significant amount of noises^4^. For example, GE might be affected by batch effect, where results obtained from two different runs of experiments may present substantial variations^5^, even when the input materials are identical. Additionally, the expression of particular genes varies with cell cycle^6^, increasing the heterogeneity observed in single cells^7^. To cope with these sources of variations, normalization of GE is usually a mandatory step before downstream functional analysis^7^. Even with these procedures, other sources of biases still exist, e.g. dependent on read depth, cell capture efficiency and experimental protocols etc.

Single nucleotide variations (SNVs) are genetic alterations of one single base occurring in specific cells as compared to the population background. SNVs may manifest their effects on gene expression by *cis* and/or *trans* effect^8,9^. The disruption of the genetic stability, e.g. increasing number of new SNVs, is known to be linked with cancer evolution^10,11^. A cell may become the precursor of a subpopulation (clone) upon gaining a set of SNVs. Considerable heterogeneity exists not only between tumors but also within the same tumor^12,13^. Therefore, investigating the patterns of SNVs provides means to understand tumor heterogeneity.

In single cells, SNVs are conventionally obtained from single-cell exome-sequencing and whole genome sequencing approaches^14^. The resulting SNVs can then be used to infer cancer cell subpopulations^15,16^. In this study, we propose to obtain useful SNV-based genetic information from scRNA-seq data, in addition to the GE information. Rather than being considered the “by-products” of scRNA-seq, the SNVs not only have the potential to improve the accuracy of identifying subpopulations compared to GE, but also offer unique opportunities to study the genetic events (genotype) associated with gene expression (phenotype)^17,18^. Moreover, when the coupled DNA- and RNA-based single-cell sequencing techniques become mature, the computational methodology proposed in this report can be adopted as well^19^.

Here we first built a computational pipeline to identify SNVs from scRNA-seq raw reads directly. We then constructed a linear modeling framework to obtain filtered, effective and expressed SNVs (eeSNVs) associated with gene expression profiles. In all the datasets tested, these eeSNVs show better accuracies at retrieving cell subpopulation identities, compared to those from gene expression (GE). Moreover, when combined with cell entities into bipartite graphs, they demonstrate improved visual representation of the cell subpopulations. We ranked eeSNVs and genes according to their overall significance in the linear models and discovered that several top-ranked genes (e.g. HLA genes) appear commonly in all cancer scRNA-seq data. In summary, we emphasize that extracting SNV from scRNA-seq analysis can successfully identify subpopulation complexity and highlight genotype-phenotype relationships.

## Results

### SNV calling from scRNA-seq data

We implemented a pipeline to identify SNVs directly from FASTQ files of scRNA-seq data, following the SNV guideline of GATK (Suppl. Figure S1). We applied this pipeline to five scRNA-seq cancer datasets (Kim^20^, Ting^21^, Miyamoto^22^, Patel^23^ and Chung^24^ see Methods), and tested the efficiency of SNV features on retrieving single cell groups of interest. These datasets vary in tissue types, origins (Mouse or Human), read lengths and map-ability (Table 1). They all have pre-defined cell types (subclasses), providing useful references for assessing the performance of a variety of clustering methods used in this study.

**Table 1:** Summary of scRNA-seq datasets used in this study.

| Data | Description | Organism | Sub- | Cell | Reads | Map- | Read | Expresse | % samples | % reads |
| --- | --- | --- | --- | --- | --- | --- | --- | --- | --- | --- |

|  |  |  | class | count | Per cell | ability | length | d genes | removed after filtering | removed after filtering |
| --- | --- | --- | --- | --- | --- | --- | --- | --- | --- | --- |
| Kim dataset<sup>20</sup> | Renal carcinoma cancer cell from patient and PDX | Human | 3 | 91 | 4.1M | 82 % | 100 | 18288 | 23 | 14 |
| Ting dataset<sup>21</sup> | Pancreas Circulating Tumor cells (CTC) Cancer | Mouse | 6 | 116 | 13.7M | 39 % | 50 | 15868 | 32 | 31 |
| Miyamoto dataset<sup>22</sup> | Prostate CTCs Cancer | Human | 24 | 133 | 2.0M | 44 % | 50 | 18224 | 22 | 40 |
| Patel dataset<sup>23</sup> | Glioblastoma tumor cells | Human | 7 | 593 | 3.2M | 63 % | 25 | 25053 | 38 | 42 |
| Chung dataset<sup>24</sup> | Breast cancer | Human | 13 | 556 | 5.9M | 81% | 100 | 22121 | 1 | 15 |

We evaluated the GATK SNV calling pipeline using several approaches. First, we estimated the true positive rates of the SNV calling pipeline at different depths of scRNA-Seq reads. For this we performed a simulation experiment by artificially introducing 50,000 random SNVs in the exonic regions of hg19 and measured recovery of these SNVs using our pipeline on Kim dataset. The true positive rates monotonically increase with read depth. For as few as 4 read-depth, the pipeline achieves on average over 50% true positive rate and increases to 68% true positive rate when read depth is more than 6 (Figure 1A). This accuracy is in line with what was reported from bulk cell RNA-Seq ^25^. The false positive rate is consistently less than 0.1, and the median reaches below 0.05 when the read depth is more than 6 (Figure 1B). We compared the SNV call results from GATK to those from another SNV caller FreeBayes^26^ and obtained similar results (Suppl. Figure S2 A and B). Additionally, we conducted simulation experiment on a new non-cancer 10X Genomic dataset, and obtained comparable true positive rates (Suppl. Figure S2 C and D). Moreover GATK shows better performance than FreeBayes in the 10X dataset. We thus opted to use GATK to call SNVs for the remainder of the report, given its popularity and performance.

**Figure 1:**
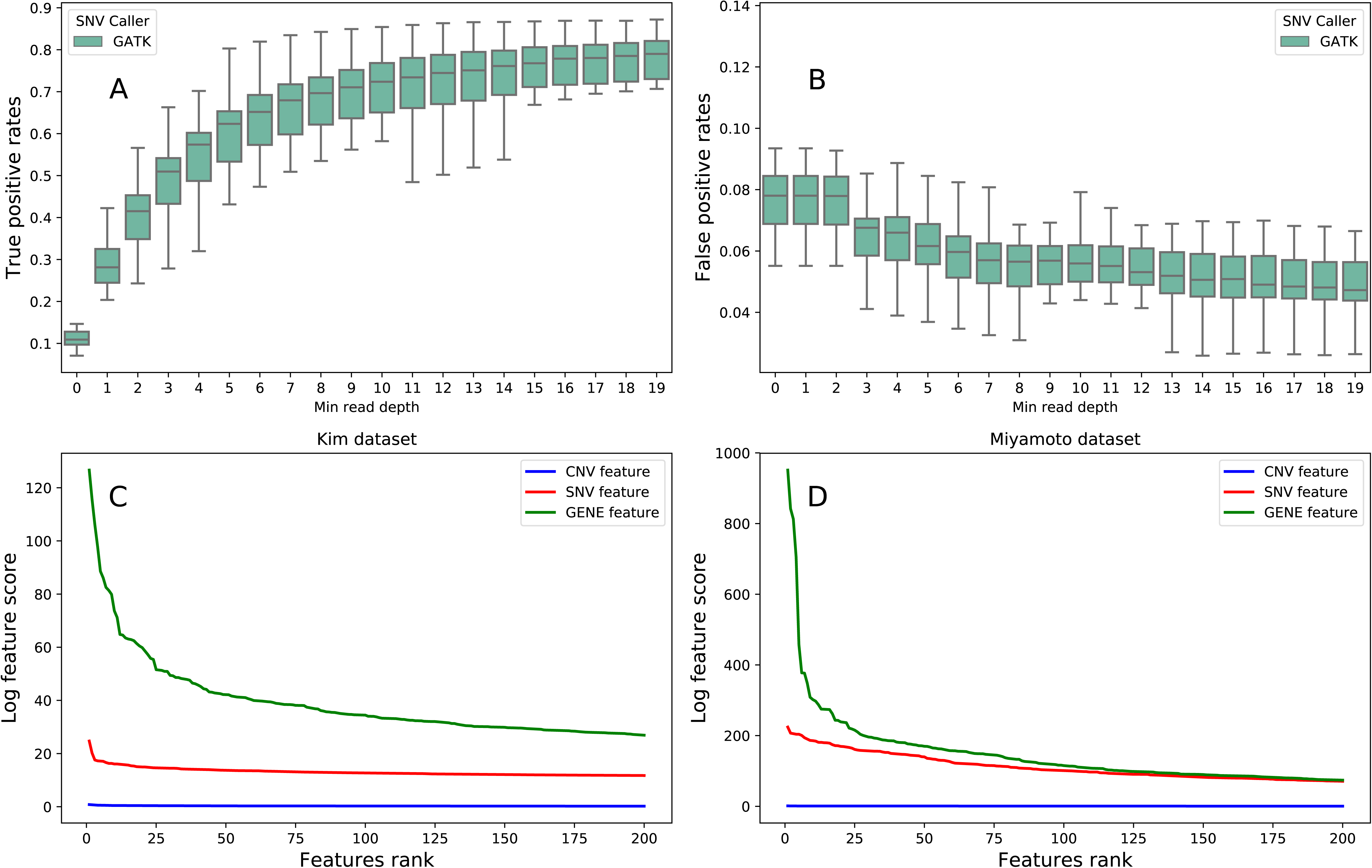
The performance measurements of GATK SNV calling and SSrGE pipelines. (A-B) Performance measurement of GATK SNV calling pipeline. Box plots of true positive rate (A) and false positive rate (B) with respect to the read depth at the called SNV position. The rates are calculated from GATK SNV calling pipeline, using hg19 reference genome to align modified scRNA-seq reads from a subset of 20 cells from the Kim dataset, which were introduced 50000 random artificial mutations the exonic region of the reads. (C-D) Comparisons of importance the different types of features in SSrGE models, with respect to the ranking, in Miyamoto dataset (C) and Kim dataset (D). The scores of the SNVs and CNVs correspond to the sum of the coefficients inferred by the SSrGE models. The gene score is the sum of the SNVs scores for a particular gene. Blue: CNV feature; Red: eeSNV feature; Green: gene feature.

### Using SSrGE to detect eeSNVs in scRNA-seq data

To link the relationship between SNV and GE, we developed a method called “Sparse SNV inference to reflect Gene Expression” (SSrGE), as detailed in Materials and Methods. In addition to SNV, we also optionally considered the effect of CNVs on gene expression, since copy number variation (CNV) may contribute to gene expression variation as well. Similar to gene-based association method PrediXscan^17^, SSrGE uses SNVs and additionally optionally CNVs as predictors to fit a linear model for gene expression, under LASSO regularization and feature selection^27^. We choose LASSO rather than elasticNet for penalization, so that the list of resulting eeSNVs is short (Suppl. Figure S3). These eeSNVs serve as refined descriptive features for subsequent subpopulation identification. To directly pinpoint the contributions of SNVs relevant to protein-coding genes, we used the SNVs residing between transcription starting and ending sites of genes as the inputs. We further assessed the relative contributions of eeSNVs and CNVs to gene expression and found that the coefficients of the CNVs are significantly lower than those of eeSNVs(Figure 1C and D). The ranks of the top genes with and without CNVs in the SSrGE models are not statistically different overall, as the Kendall-Tau correlation scores^28^ are close to 1 with p-values=0.

Additionally, SNV genotypes and allelic specific level gene expression may also complicate the relationships between eeSNVs and gene expression. Therefore, we further calibrated SSrGE model by considering SNV genotype and allelic specific gene expression. We used QUASAR^29^ to estimate the SNV genotypes (Suppl. Table 1), and the allelic specific gene expression using the SNV genotype. We rebuilt individual SSrGE models using only the SNVs from a particular genotype and allelic-specific gene expression, and then merged the eeSNV weights from related SSrGE models together to obtain a final ranking of eeSNVs. The new rankings are not statistically different compared to the previous approach (Suppl. Table 1). The Kendall Tau scores, which evaluate the similarities between the re-calibrated model and the original model have p-values =0 in all data sets.

Lastly, to quantitatively evaluate if the eeSNVs obtained from SSrGE are truly significant, we designed a simulation pipeline (Methods). The pipeline creates random binary matrices of SNVs for *n* simulated cells, which are connected to the matrices of gene expression. The SNVs present in the simulated cell have probabilities to modify gene expression of the genes positively or negatively. We used various levels of noise to perturb the GE and the SNV matrices. We compared the ranks of top genes identified by SSrGE to the expected impact of each gene provided by the simulation. The inferred top ranked genes using SSrGE have monotonic and positive correlations with those set by the simulation (Suppl. Figure S4A). These correlations are all significant (p-value << 0.05), independently of the alpha and the level of noise used, confirming the value of SSrGE model. Moreover, to simulate the patterns of dropout in the data, we also introduced two other parameters, one for random dropout or biased dropout towards both lowly expressed genes, and other for dropout rate relative to cell, gene or reads (Methods). We observed that SSrGE performs well on all dropout models (Suppl. Figure S4B). Thus, SSrGE method is validated to generate eeSNVs that are truly important.

### eeSNVs are better than gene expression at identifying subpopulations

We measured the performance of SNVs and gene expression (GE) to identify subpopulations on the five datasets, using five clustering approaches (Figure 2). These clustering approaches include two dimension reduction methods, namely Principal Component Analysis (PCA)^30^ and Factor Analysis (FA)^31^, followed by either K-Means or the hierarchical agglomerative method (agglo) with WARD linkage^32^. We also used a recent algorithm SIMLR designed explicitly for scRNA-seq data clustering and visualization ^33^. To evaluate the accuracy of obtained subpopulations in each dataset, we used the metric of Adjusted Mutual Information (AMI) over 30 bootstrap runs, from the optimal *a* parameters (Suppl. Table S2). These optimal parameters were estimated by testing different *a* values for each dataset and each clustering approach (Suppl. Figure S5). As shown in Figure 2, eeSNVs are better features to retrieve cancer cell subpopulations compared to GE, independent of the clustering methods used. Among the clustering algorithms, SIMLR tends to be a better choice using eeSNV features.

**Figure 2:**
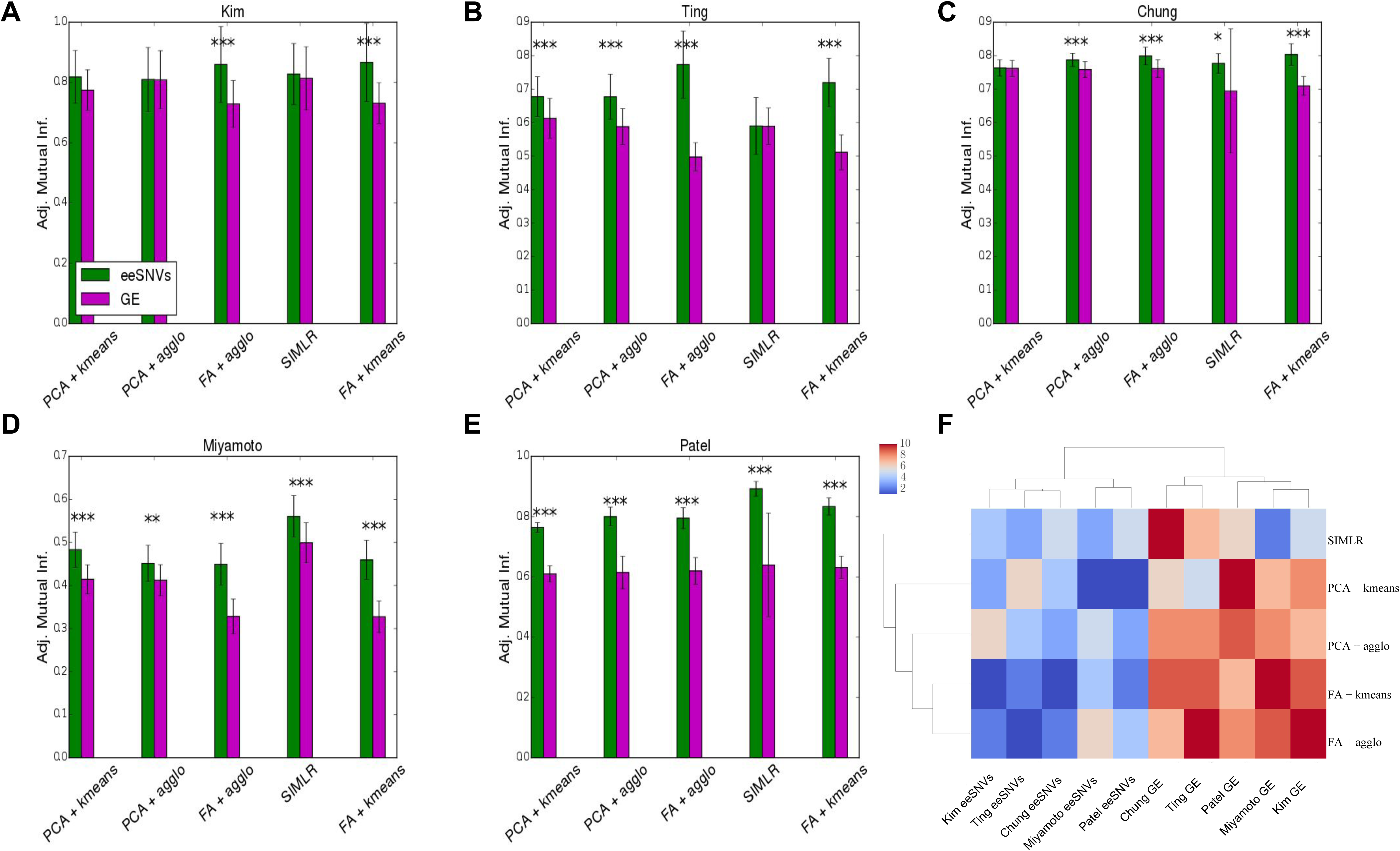
Comparison of clustering accuracy using eeSNV and gene expression (GE) features. (A-E) Bar plots comparing the clustering performance using eeSNV vs. gene expression (GE) as features, over five different clustering strategies and five datasets, each with its own pre-defined classes as the truth measure: (A) Kim dataset (B) Ting dataset (C) Chung dataset (D) Miyamoto dataset and (E) Patel dataset. Y-axis is the adjusted mutual information (AMI) obtained across 30 bootstrap runs (mean ± s.d.). *: P<0.05, ** P<0.01 and *** P<0.001. (F) Heatmap of the rankings among different methods and datasets as shown in (A-E).

Additionally, we also computed the Adjusted Rand index (ARI)^34^ and V-measure^35^, two other metrics for modularity measurements (Methods) and obtained similar trends (Suppl. Figure S6). Similar to AMI, ARI is a normalized metric against random chance, and evaluates the number of correct pairs obtained. On the other hand, V-measure combines the homogeneity score, which measures the homogeneity of reference classes in the obtained clusters, and the completeness score, which measures the homogeneity of obtained clusters within the reference classes. Due to the high number of small homogenous clusters obtained for the Miyamoto dataset, we observed higher V-measure scores, compared to AMI and ARI results (Suppl. Figure S6).

### Visualization of subpopulations with bipartite graphs

Bipartite graphs are an efficient way to describe the binary relations between two different classes of objects. We next represented the presence of the eeSNVs into single cell genomes with bipartite graphs using ForceAtlas2 algorithm^36^. We drew an edge (link) between a cell node and a given eeSNV node whenever an eeSNV is detected. The results show that bipartite graph is a robust and more discriminative alternative (Figure 3), comparing to PCA plots (using GE and eeSNVs) as well as SIMLR (using GE). For Kim dataset, bipartite graph separates the three classes perfectly. However, gene-based visualization approaches using either PCA or SIMLR have misclassified data points. For Ting data, the eeSNV-cell bipartite graph gives a clear visualization of all six different subgroups of single cells. Other three approaches have more exaggerated separations among the same mouse circulating tumor cells (CTC) subgroup MP (orange color), but mix some other subpopulations (e.g. GM, MP and TuGMP groups). Miyamoto dataset is the most difficult one to visualize among the four datasets, due to its high number (24) of reference classes and heterogeneity among CTCs. Bipartite graphs are not only able to condense the whole populations, but also separate subpopulations (e.g. the orange colored PC subpopulation) much better than the other three methods.

**Figure 3:**
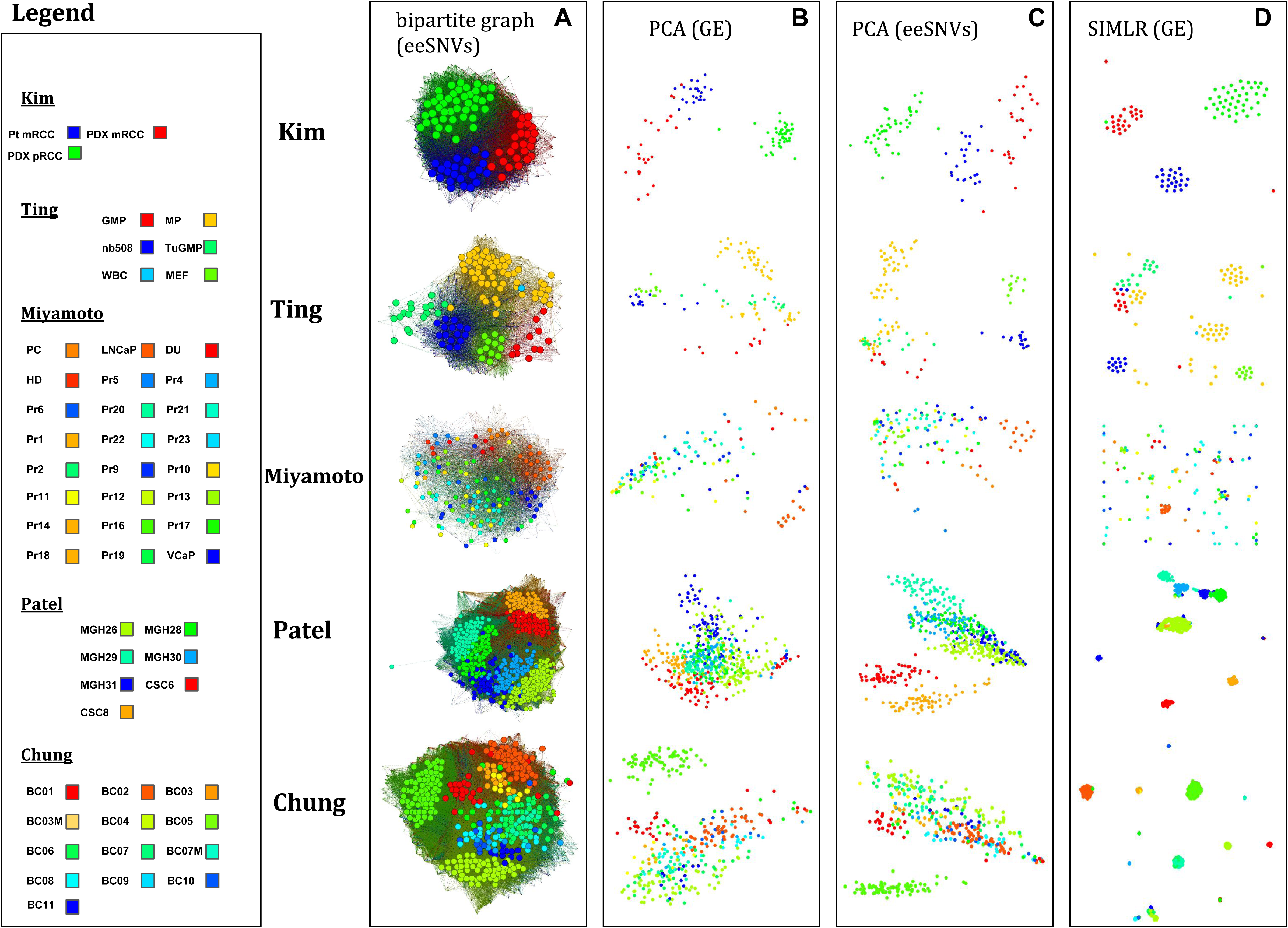
Comparison of clustering visualization using eeSNV and gene expression (GE) features. (A) Bipartite graphs using eeSNVs and cells as two groups of nodes. An edge between a cell and an eeSNV represents the presence of the eeSNV within that cell. (B) Principle Component Analysis (PCA) results using GE as features of the cells. (C) PCA results using eeSNVs as features of the cells. (D) SIMILR results using GE as the input.

### Characteristics of eeSNVs

In SSrGE, regularization parameter *a* is the only tuning variable, controlling the sparsity of the linear models and influences the number of eeSNVs. We next explored the relationship between eeSNVs and *a* (Figure 4). For every dataset, increasing the value of *a* decreases the number of selected eeSNVs overall (Figure 4A), as well as the average number of eeSNVs associated with every expressed gene (Figure 4B). The optimal *a* depends on the clustering algorithm and the dataset used (Suppl. Table S2 and Suppl. Figure S5). Increasing the value of *a* expands the proportion of eeSNVs that have annotations in human dbSNP138 database, indicating a higher true positive rate of SNVs compared to that prior to SSrGE filtering (Figure 4C). Additionally, increasing *a* increases the average number of cells sharing the same eeSNVs (Figure 4D), corresponding to the decreasing number of eeSNVs (Figure 4B). Note the slight drop in the average number of cells sharing the same eeSNVs in Kim data when *a* > 0.6, this is due to over-penalization (eg. *a* =0.8 yields only 34 eeSNVs).

**Figure 4:**
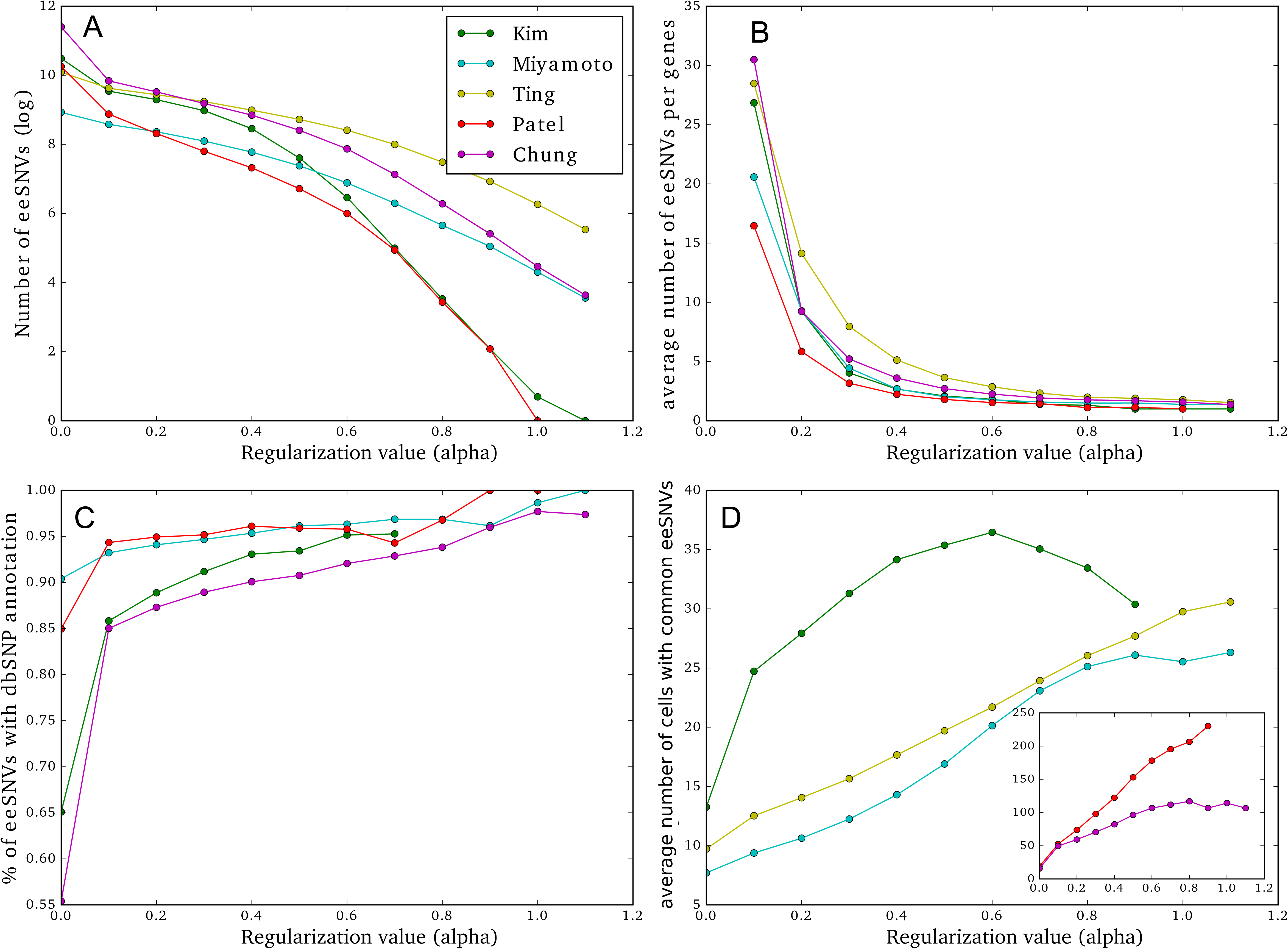
Characteristics of the eeSNVs. X-axis: the regularization parameter *a* values used by LASSO penalization in the SSrGE models. And the Y-axes are: (A) Log10 transformation of the number of eeSNVs. (B) The average number of eeSNVs per gene. (C) The proportion of SNVs with dbSNP138 annotations (human datasets). (D) The average number of cells sharing eeSNVs.

### Cancer relevance of eeSNVs

Following the simulation results, we ranked the different eeSNVs and the genes for the five datasets, from SSrGE models (Suppl. Tables S3). We found that eeSNVs from multiple genes in Human Leukocyte Antigen (HLA) complex, such as HLA-A, HLA-B, HLA-C and HLA-DRA, are top ranked in all four human datasets (Table 2 and Suppl. Tables S3). HLA is a family encoding the major histocompatibility complex (MHC) proteins in human. Beta-2-microglobulin (B2M), on the other hand, is ranked 7^th^ and 45^th^ in Ting and Patel datasets, respectively (Table 2). Unlike HLA that is present in human only, B2M encodes a serum protein involved in the histocompatibility complex MHC that is also present in mice. Other previously identified tumor driver genes are also ranked top by SSrGE, demonstrating the impact of mutations on cis-gene expression (Table 2 and Suppl. Tables S3). Notably, *KRAS*, previously linked to tumor heterogeneity (Kim et al., 2015), is ranked 13^th^ among all eeSNV containing genes (Suppl. Tables S3). *AR* and *KLK3*, two genes reported to show genomic heterogeneity in tumor development in the original study^22^, are ranked 6^th^ and 19^th^, respectively. *EGFR*, the therapeutic target in Patel study with an important oncogenic variant EGFRvIII (Patel et al., 2014), is ranked 88^th^ out of 4,225 genes. Therefore, genes top-ranked by their eeSNVs are empirically validated.

**Table 2:** A list of interested genes highly ranked. Ranks with ‘*’ designate cancer driver genes reported in the original studies.

| **Dataset** | **Kim** | **Patel** | **Miyamoto** | **Chung** | **Ting (mouse)** |
| --- | --- | --- | --- | --- | --- |
| *HLA-A* | **32** | **8** | **2** | **6** | - |
| *HLA-B* | **3** | **105** | **1** | **4** | - |
| *HLA-C* | **1** | **98** | **4** | **2** | - |
| *HLA-DRA* | **71** | 771 | 200 | **18** | - |
| *B2M* | 1617 | **45** | 301 | 425 | **7** |
| *KRAS* | **13** | 2101 | 2254 | **72** | 235* |
| *TRP53* | NA | NA | NA | NA | 365* |
| *SPARC* | **22** | **37** | 567 | 2070 | **79** |
| *EGFR* | 2231 | **88\*** | NA | NA | NA |
| *AR* | NA | NA | **6\*** | 340 | NA |
| *KLK3* | NA | NA | **19\*** | NA | NA |

Next we conducted more systematic investigation to identify KEGG pathways enriched in each dataset, using these genes as the input for DAVID annotation tool^37^ (Figure 5A). The pathway-gene bipartite graph illustrates the relationships between these genes and enriched pathways (Figure 5B). As expected, Antigen processing and presentation pathway stands out as the most enriched pathway, with the sum −log10 (p-value) of 15.80 (Figure 5B). “Phagosome” is the second most enriched pathway in all four data sets, largely due its members in HLA families (Figure 5B). Additionally, pathways related to cell junctions and adhesion (focal adhesion and cell adhesion molecules CAMs), protein processing (protein processing in endoplasmic reticulum and proteasome), and PI3K-AKT signaling pathway are also highly enriched with eeSNVs (Figure 5A).

**Figure 5:**
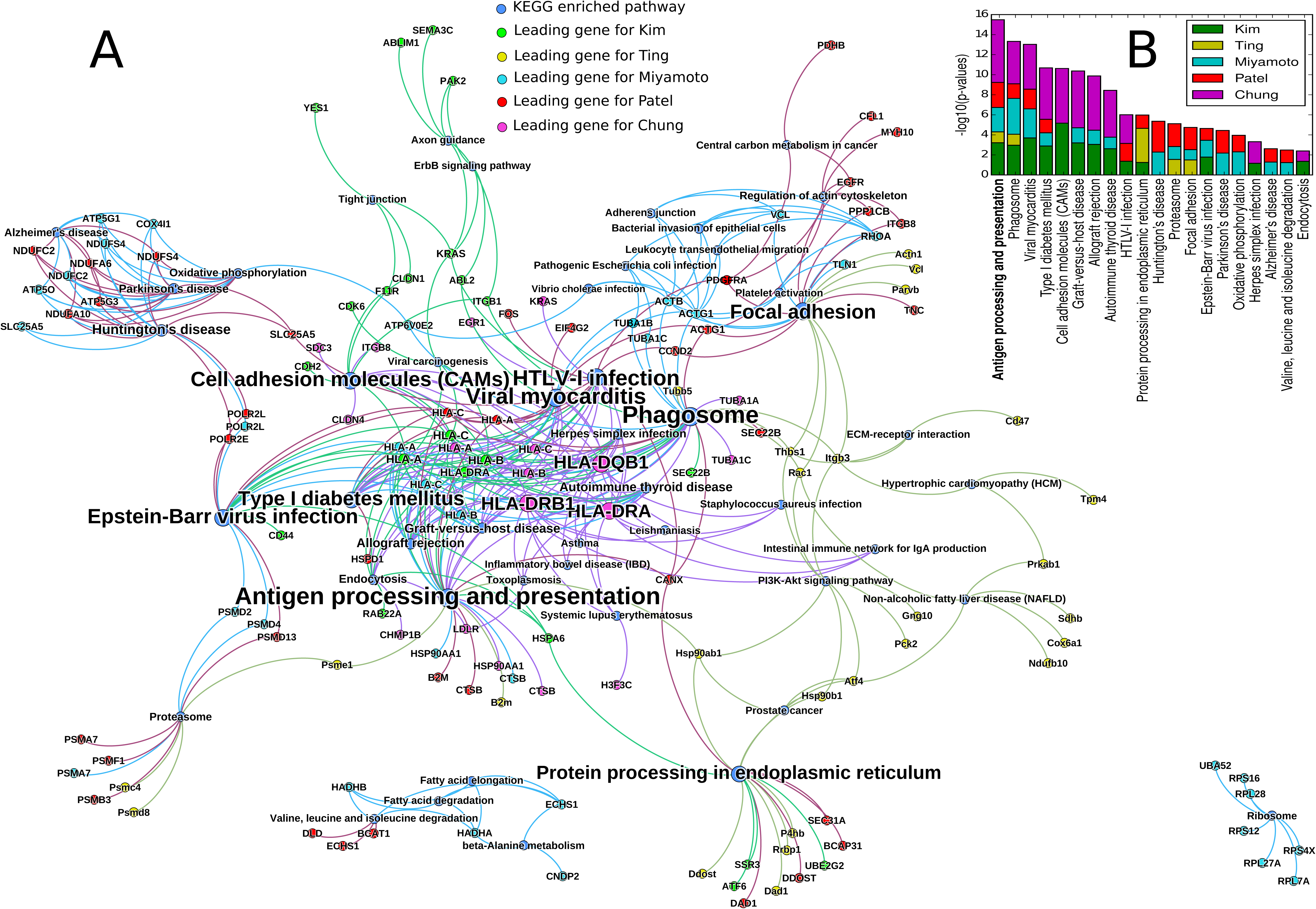
Gene and KEGG pathways enriched with eeSNVs in the five scRNA-seq datasets. (A) Bipartite graph using significant KEGG pathways and genes enriched with eeSNVs as nodes. An edge exists between a significant pathway and a gene if this gene is part of the pathway. Genes of each data set is represented with a distinct color. The size of the nodes reflects the gene and the pathway scores. The gene scores are computed by SSrGE and the pathway scores are the sum of the gene scores linked for each pathway. (B) KEGG pathways enriched within the top 100 genes based on eeSNV contributions in the <u>five</u> datasets. Pathways are sorted by the sum of the −log10 (p-value) of each dataset, in the descending order.

### Heterogeneity markers using eeSNVs

We display the potential of eeSNV as heterogeneity markers via pseudo-time reconstruction and hearmap, using Kim dataset (Figure 6A and B). We built a Minimum Spanning Tree, similarly to the Monocle algorithm^38^, to reconstruct the pseudo-time ordering of the single-cells. The graphs beautifully capture the continuity among cells, from the primary to metastasized tumors (Figure 6A). Moreover, it highlights ramifications inside each of the subgroups, demonstrating the intra-group heterogeneity. On the contrary, pseudo-time reconstruction using GE showed much less complexity and more singularity (Supplementary Figure S7). As a confirmation, hierarchical clustering of eeSNV heatmp also shows almost perfect separation of the three subgroups (Figure 6B). Next, we used our method to identify eeSNVs specific to each single-cell subgroup and ranked the genes according to these eeSNVs. We compared the characteristics of the metastasis cells to primary tumor cells. Two top-ranked genes identified by the method, CD44 (1^th^) and LPP (2^th^), are known to promote cancer cell dissemination and metastasis growth after genomic alteration^39–42^ (Suppl. Table S3). Other top-ranked genes related to metastasis are also identified, including LAMPC2 (7^th^), HSP90B1 (14^th^), MET (44^th^) and FN1 (52^th^). As expected, “Pathways in Cancer” are the top-ranked pathway enriched with mutations (Figure 6B). Additionally, “Focal Adhesion”, “Endocytosis” pathways are among the other significantly mutated pathways, providing new insights on the mechanistic difference between primary and metastasized RCC tumors (Figure 6C).

**Figure 6:**
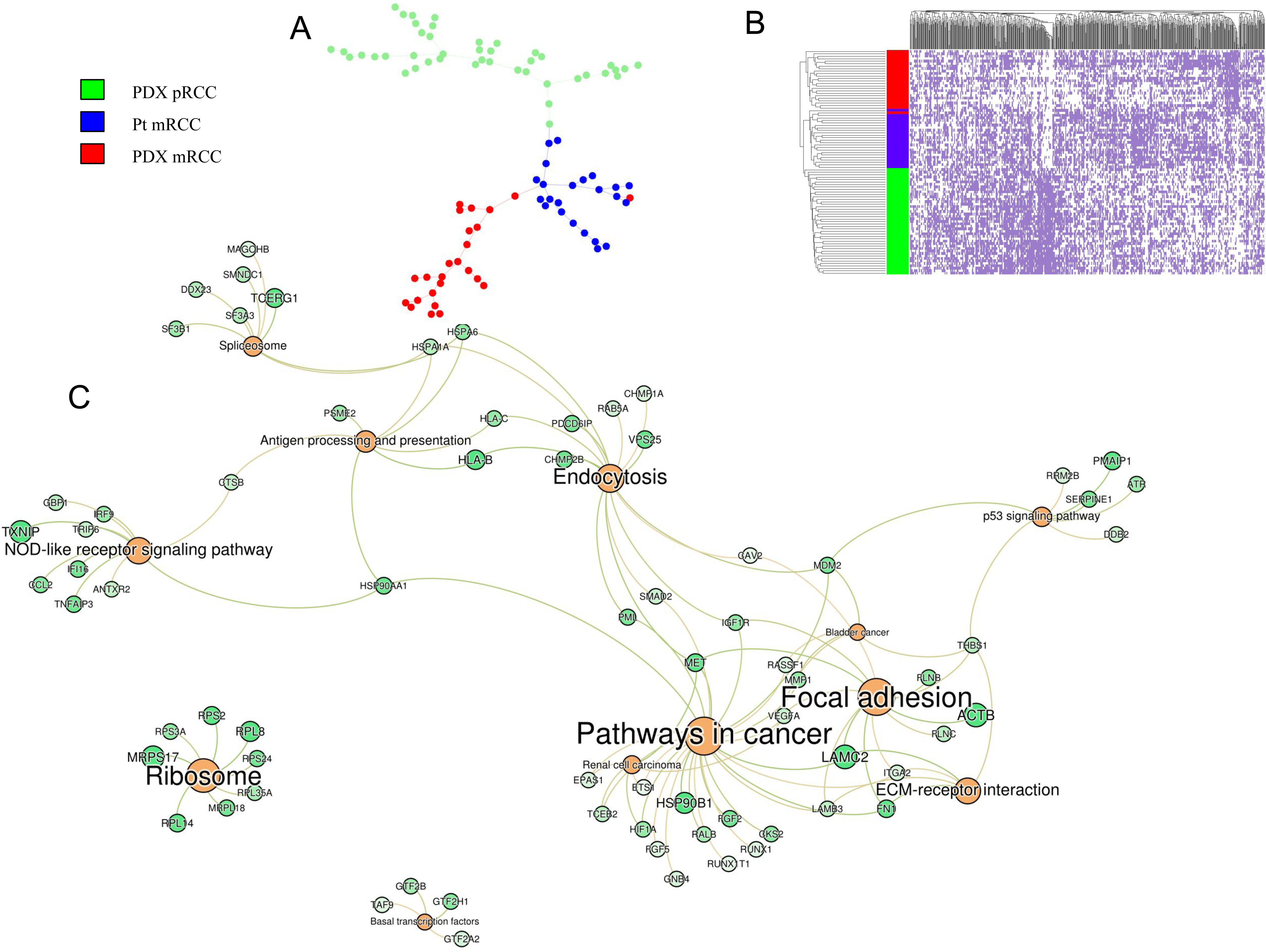
Heterogeneity revealed by Kim dataset. (A) Pseudo-time ordering reconstruction of the different subgroups: single-cells from PDX primary tumor (green), Patient metastasis (blue) and PDX metastatis (red). The eeSNVs are obtained with *a* = 0.6. The tree is inferred using the MST algorithm on Pearson’s correlation-based distance matrix. (B) Heatmap of the cells (row) and eeSNVs (column). (C) Bipartite graph using KEGG pathways and genes enriched with significant eeSNVs as two set of nodes. The significant eeSNVs are inferred from the metastasized cells, compared to the primary tumor cells. The size of the nodes reflects the gene scores (given by SSrGE) and the pathway scores (sum of the gene scores). Lighter green indicates genes with a lower rank.

Another application is to explore the potential of eeSNVs to separate different cell types within the same individual. Towards this, we extended the same analysis on the two patients BC03 and BC07 from the Chung dataset, who have primary and metastasized tumor cells as well as infiltrating immune cells. Again, bipartite graphs and minimum spanning trees-based visualization illustrate clear separations of tumor (primary and metastasized) cells from immune cells (Suppl. Figure S8). Furthermore, the top-ranked genes relative to the metastasis subgroups (BC03M and BC07M) present some similarities with those found in Kim dataset (Suppl. Tables S4). Strikingly, CD44 is also top-ranked (23^th^) among the significant genes of BC07M. Similarly, HSP90B1 is top-ranked as the 63^th^ and 51^th^ most important genes, in BC03M and BC07M respectively.

### Integrating DNA- and RNA-Seq data measured in the same single cells

Coupled DNA-Seq and RNA-Seq measurements from the same single cell are the new horizon of single-cell genomics. To demonstrate the potential of SSrGE in integrating DNA and RNA data, we downloaded a public single cells data, which have DNA methylation and RNA-Seq records from the same hepatocellular carcinoma (HCC) single cells (Hou dataset)^43^. We then inferred SNVs from the aligned reduced representation bisulfite sequencing (RRBS) reads (see Methods), and used them to predict the scRNA-Seq data from the same samples. Given the fact that SNVs are heterozygous among tumor and normal cells, and that a small fraction of genes harboring eeSNVs are subject to CNV, we included both the percentages of SNVs as well as CNVs as additional predictive variables in the SSrGE model besides SNV features. Interestingly, the identified eeSNVs can clearly separate normal hepatocellular cells from cancer cells and highlight the two cancer subtypes identified in the original study (Figure 7). Pseudo-time ordering shows an early divergence between the two previously assumed subtypes (Figure 7B). This observation is confirmed by hierarchical clustering of eeSNV based heatmap (Figure 7C). A simplified version of SSrGE model, where only SNV features were considered as predictors for gene expression, shared 92% eeSNVs as those in Figure 7A, and achieved almost identical separations between normal hepatocellular cells and cancer cells (data not shown). This confirms the earlier observation that eeSNVs are much more important predictive features, compared to CNVs (Figure 1C and D).

**Figure 7:**
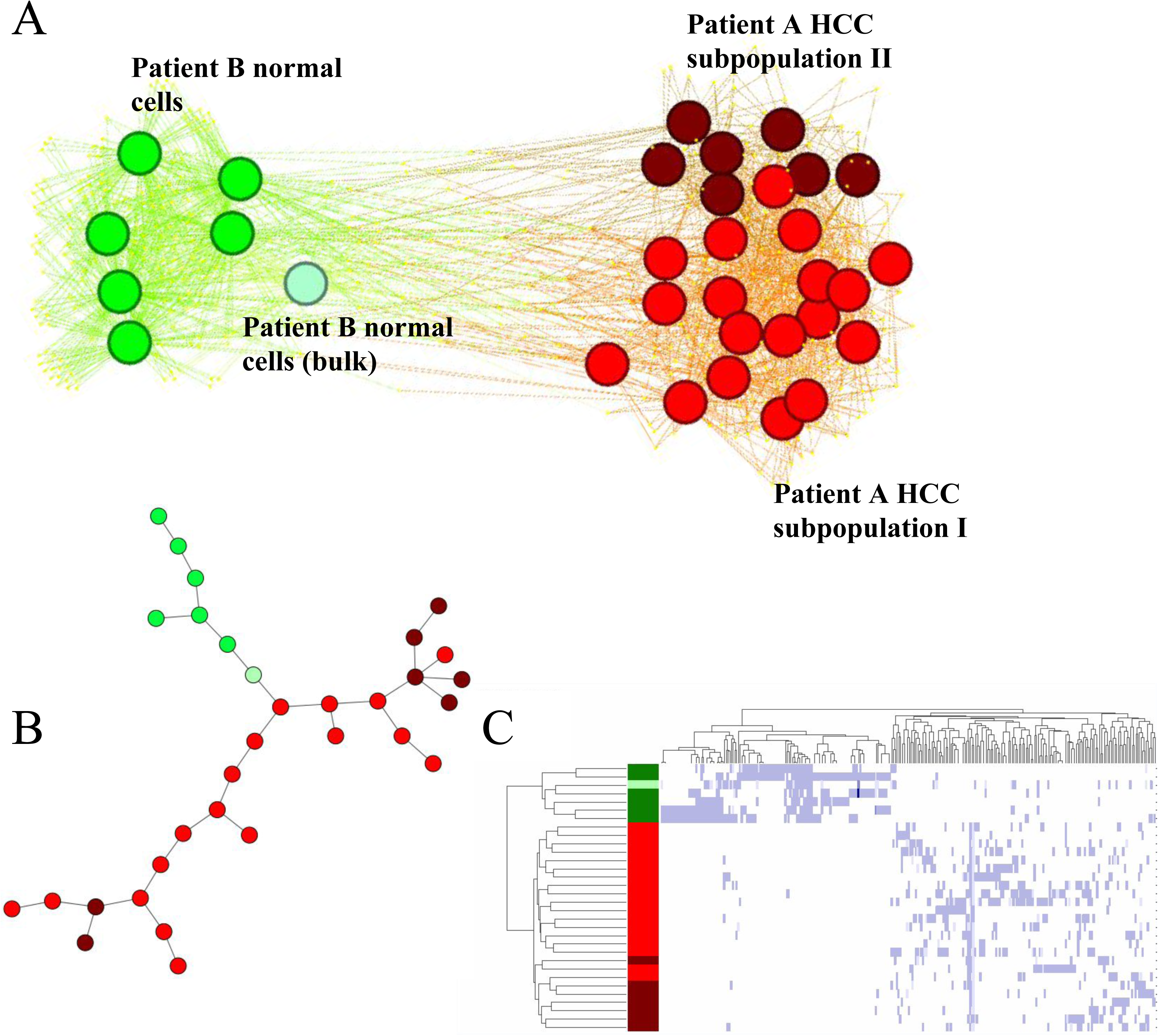
Heterogeneity revealed by eeSNVs from multi-omics single cell HCC (Hou) dataset. Normal cells are colored green and HCC tumor cells are colored dark or bright red. (A) Bipartite-graph representation using the single cells and eeSNVs from RRBS reads as two sets of nodes. (B) Pseudo-time ordering reconstruction of the HCC cells, using eeSNVs in from RRBS. (C) Heatmap of the cells (row) and eeSNVs (column).

We postulated that a considerable part of bisulfite reads was aligned with methylation islands associated with gene promoter regions. Thus, we annotated eeSNVs within 1500bp upstream of the transcription starting codon, and obtained genes with these eeSNVs, which are significantly prevalent in certain groups. When comparing HCC vs. normal control cells, two genes PRMT2, SULF2 show statistically significant mutations in HCC cells (P-values < 0.05). Down-regulation of PRMT2 was previously associated with breast cancer^44^, SULF2 was known to be up-regulated in HCC and promotes HCC growth^45^.

## Discussion

Using GE to accurately analyze scRNA-seq data has many challenges, including technological biases such as the choice of the sequencing platforms, the experimental protocols and conditions. These biases may lead to various confounding factors in interpreting GE data^5^. SNVs, on the other hand, are less prone to these issues given their binary nature. In this report, we demonstrate that eeSNVs extracted from scRNA-seq data are ideal features to characterize cell subpopulations. Moreover, they provide a means to examine the relationship between eeSNVs and gene expression in the same scRNA-seq sample.

### eeSNVs have improved accuracy in identifying single-cell subpopulations

The process of selecting eeSNVs linked to GE allows us to identify representative genotype markers for cell subpopulations. We speculate the following reasons attributed to the better accuracies of eeSNVs compared to GE. First, eeSNVs are binary features rather than continuous features like GE. Thus, eeSNVs are more robust at separating subpopulations. We have noticed that SNVs are less affected by batch effects (Suppl. Figure S9). Secondly, LASSO penalization works as a feature selection method and minimizes the spurious SNVs (false positive) from the filtered set of eeSNVs. Thirdly, since eeSNVs are obtained from the same samples as scRNA-seq data, they are more likely to have biological impacts, and this is supported the observation that they have high prevalence of dbSNP annotations.

A small number of eeSNVs can be used to discriminate distinct single-cell subpopulations, as compared to thousands of genes that are normally used for scRNA-seq analyses. Taking advantage of the eeSNV-GE relationship, a very small number of top eeSNVs still can clearly separate cell subpopulations of the different datasets (eg. 8 eeSNV features have decent separations for Kim dataset). Moreover, our SSrGE package can be easily parallelized and process each gene independently. It has the potential to scale up to very large datasets, well-poised for the new wave of scRNA-seq technologies that can generate thousands of cells at one time^46^. One can also easily rank the eeSNVs and the genes harboring them, for the purpose of identifying robust eeSNVs as genetic markers for a variety of cancers.

The eeSNV based approach to identify subpopulations can be generalized to other non-cancer tissues and scRNA-Seq data generated from various platforms. Additionally we analyzed 9 datasets, including two non-cancer 10X Genomics datasets with a large number of cells, four non-cancer datasets (GSE79457, GSE81903, GSE71485, and GSE80232), and three cancer single cell datasets (GSE69405, GSE81383, and GSE65253). Bipartite graphs with eeSNVs and cells were able to separate the single-cell subpopulations based on their genotype (Suppl. Figure S10A-I).

### eeSNVs highlight genes linked to cancer phenotypes

SSrGE uses an accumulative ranking approach to select eeSNVs linked to the expression of a particular gene. Mainly, HLA class I genes (HLA-A, HLA-B and HLA-C) are top-ranked for the three human datasets, and they contribute to “antigen processing and presentation pathway”, the most enriched pathways of the four datasets. HLA has amongst the highest polymorphic genes of the human genome^47^, and the somatic mutations of genes in this family occurred in the development and progression of various cancers^48,49^. eeSNVs of HLA genes could be used as fingerprints to identify the cellular state of the cancer cells, and lead to better separation between the primary (pRCC, green) and metastatic cells (mRCC, red and blue) compared to GE of HLA genes (Suppl. Figure 11). *B2M*, another gene with top-scored eeSNVs in Ting and Patel datasets, is also known to be a mutational hotspot^50^. It is immediately linked to immune response as tumor cell proliferation^48,50^. Many other top-ranked genes, such as *SPARC*, were reported to be driver genes in the original studies of the different dataset. Thus, it is reasonable to speculate that SSrGE is capable of identifying some driver genes. Another possibility is that some of the eeSNVs reflect aberrant splicing of genes such as the HLA family, which are regularly found in deregulated cancer cells^51^. Nevertheless, SSrGE may miss some driver mutations due to the incomplete DNA coverage due to the use of scRNA-Seq reads. Also, its primary goal is to identify a minimal set of eeSNV features by LASSO penalization but in case of correlated features, LASSO may select one of those highly correlated SNV features that correspond to GE.

### eeSNVs reveal higher degree of single-cell heterogeneity than gene expression

We have showed with strong evidence that eeSNVs unveil inter- and intra-tumor cells heterogeneities better than gene expression count data obtained from the same RNA-Seq reads. Reconstructing the pseudo-time ordering of cancer cells from the same tumor (Kim dataset) displays branching even inside primary tumor and metastasis subgroups, which gene expression data are unable to do. We identified genes enriched with SNVs specific to the metastasis, which were not reported in the original HCC single cell study^43^. Most interestingly, we showed that eeSNVs can also be retrieved from RRBS reads in a multi-omics single-cell HCC dataset, a twist from their original purpose of single-cell DNA methylation. Again, genes ranked by eeSNVs from RRBS reads only differentiate normal from cancer cells but also the different cancer subtypes. We identified several genes that are significant in either HCC or HCC subgroup, whose promoters are highly impacted by eeSNVs. Thus, we have demonstrated that our method is on the fore-front to analyze data generated by new single-cell technologies extracting multi-omics from the same cells^43,52^.

### Advantages of using bipartite graphs to represent scRNA-seq data

Bipartite graphs are a natural way to visualize eeSNV-cell relationships. We have used force-directed graph drawing algorithms involve spring-like attractive forces and electrical repulsions between nodes that are connected by edges. This approach has the advantage to reveal “outlier” single cells, with a small set of eeSNVs, compared to those distance-based approaches. Moreover, the bipartite representation also reveals directly the relationship between single cells and the eeSNV features. Contrary to dimension reduction approaches such as PCA that requires linear transformation of features into principle components, bipartite graphs preserve all the binary information between cell and eeSNV. Graph analysis software such as Gephi^36^ or Cytoscape^53^ can be utilized to explore the bipartite relationships in an interactive manner.

## Conclusion

We demonstrated the efficiency of using eeSNVs for cell subpopulation identification over multiple datasets. eeSNVs are excellent genetic markers for intra-tumor heterogeneity and may serve as genetic candidates of new treatment options. We also have developed SSrGE, a linear model framework that correlates genotype (eeSNV) and phenotype (GE) information in scRNA-seq data. Moreover, we have showed the capacity of SSrGE in analyzing multi-omics data from the same single cells, obtained from the most cutting-edge genomics techniques^54,55^. Our method has the great promise as part of routine analyses in scRNA-seq pipelines^56^, as well as multi-omics single-cell integration projects.

## Materials and Methods

### scRNA-seq datasets

All five datasets were downloaded from the NCBI Gene Expression Omnibus (GEO) portal^57^.

*Kim dataset* (*accession GSE73121*): contains three cell populations from matched primary and metastasis tumor from the same patient^20^. Patient Derivated Xenographs (PDX) were constructed using cells from the primary Clear Cell Renal Cell Carcinoma (PDX-pRCC) tumor and from the lung metastasic tumor (PDX-mRCC). Also, metastatic cells from the patient (Pt-mRCC) were sequenced.

*Patel dataset* (*accession GSE57872*): contains five glioblastoma cell populations isolated from 5 individual tumors from different patients (MGH26, MGH28, MGH29 MGH30 and MGH31) and two gliomasphere cell lines, CSC6 and CSC8, used as control^23^.

*Miyamoto dataset* (*accession GSE67980*): contains 122 CTCs from Prostate cancer from 18 patients, 30 single cells derived from 4 different cancer cell lines: VCaP, LNCaP, PC3 and DU145, and 5 leukocyte cells from a healthy patient (HD1)^22^. A total of 23 classes (18 CTC classes + 4 cancer cell lines + 1 healthy leukocyte cell lines) were obtained.

*Ting dataset* (*subset of accession GSE51372*): contains 75 CTCs from Pancreatic cancer from 5 different KPC mice (MP2, MP3, MP4, MP6, MP7), 18 CTCs from two GFP-lineage traced mice (GMP1 and GMP2), 20 single cells from one GFP-lineage traced mouse (TuGMP3), 12 single cells from a mouse embryonic fibroblast cell line (MEF), 12 single cells from mouse white blood (WBC) and 16 single cells from the nb508 mouse pancreatic cell line (nb508)^21^. KPC mice have uniform genetic cancer drivers (Tp53, Kras). Due to their shared genotype, we merged all the KPC CTCs into one single reference class. CTCs from GMP1 did not pass the QC test and were dismissed. CTCs from GMP2 mice were labeled as GMP. Finally, 6 reference classes were used: MP, nb508, GMP, TuGMP, MEF and WBC.

*Hou dataset* (*accession GSE65364*): contains 25 hepatocellular carcinoma single-cells (Ca) extracted from the same patient and 6 normal liver cells (HepG2) obtained from the adjacent normal tissue of another HCC patient^43^. The 32 cells were sequenced using scTrio-seq in order to obtain reads from both RNA-seq and reduced representation bisulfate sequencing (RRBS). The authors highlighted that one of the Ca cells (Ca_26) was likely to be a normal cell, based on CNV measurements, and thus we discarded this cell. We used the RRBS reads to infer the SNVs. We use gene expression data provided by the authors to construct a GE matrix. For controls, we used the bulk genome of all the RNA-Seq and RRBS reads of the HepG2 group.

*Chung dataset* (*accession GSE75688*): contains 549 single cells from primary breast cancer and lymph nodes metastases, extracted from 11 patients (BC01-11) of distinct molecular subtypes. BC01-02 are estrogen receptor positive (ER+); BC04-06 are human epidermal growth factor receptor 2 positive (HER+); BC03 is double positive (ER+ and HER+); BC07-11 are triple negative breast cancer (TNBC)^24^. Only BC03 and BC07 presented cells extracted from lymph nodes metastases (BC03M and BC07M). Additionally, the dataset contains a large part of infiltrating tumor cells. Following the original analytical procedure of the original study^24^, we performed an unsupervised clustering analysis to separate the cancer from the immune cells. We first reduced the dimension with a PCA analysis and then used a Gaussian mixture model to infer the clusters. We obtained a total of 372 cancer cells and 177 immune cells.

### SNV detection using scRNA-seq data

The SNV detection pipeline using scRNA-seq data follows the guidelines of GATK (http://gatkforums.broadinstitute.org/wdl/discussion/3891/calling-variants-in-rnaseq). It includes four steps: alignment of spliced transcripts to the reference genome (hg19 or mm10), BAM file preprocessing, read realignment and recalibration, and variant calling and filtering (Suppl. Figure S1)^58^.

Specifically, FASTQ files were first aligned using STAR aligner^59^, using mm10 and hg19 as reference genomes for mouse and human datasets, respectively. The BAM file quality check was done by FastQC^60^, and samples with lower than 50% of unique sequences were removed (default of FastQC). Also, samples with more than 20% of the duplicated reads were removed by STAR. Finally, samples with insufficient reads were also removed, if their reads were below the mean minus two times the standard deviation of the entire single-cell population. The summary of samples and reads filtered by these steps is listed in Table 1. Raw gene counts *X*_*j*_ were estimated using featureCounts^61^, and normalized using the logarithmic transformation:

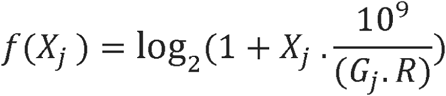

where *X*_*j*_ is the raw expression of gene j, *R* is the total number of reads and *G*_*j*_ is the length of the gene j. BAM files were pre-processed and reordered using Picard Tools (http://broadinstitute.github.io/picard/), before subject to realignment and recalibration using GATK tools^62^. SNVs are then calculated and filtered using the GATK tools *HaplotypeCaller* and *VariantFiltration* using default parameters.

Additionally, we used Freebayes^26^ with default parameters to infer the SNVs, as alternative to the *HaplotypeCaller* and *VariantFiltration* softwares. The SNV calling results between the two callers are very similar (Suppl. Figure 2).

### SNV detection using RRBS data

We first aligned the RRBS reads on the hg19 reference genome using the Bismark software ^63^. We then processed the bam files using all the preprocessing steps as described in “SNV detection using scRNA-seq data” section (i.e. Picard Preprocessing, Order reads, Split reads and Realignments), except the base recalibration step. Finally, we called the SNVs using the BS-SNPer software (default setting)^64^. The details are the following.

--minhetfreq 0.1 # Threshold of frequency for calling heterozygous SNP

--minhomfreq 0.85 # Threshold of frequency for calling homozygous SNP

--minquali 15 # Threshold of base quality

--mincover 10 # Threshold of minimum depth of covered reads

--maxcover 1000 # Threshold of maximum depth of covered reads

--minread2 2 # Minimum mutation reads number

--errorate 0.02 # Minimum mutation rate

--mapvalue 20 # Minimum read mapping value

### 10X Genomic data preprocessing

We downloaded the fastq files of two data sets: bone marrow mononuclear cells (BMMC) of 2 individuals and peripheral blood mononuclear cells (PBMC), from the 10X Genomic website (https://support.10xgenomics.com/single-cell-gene-expression/datasets). We de-multiplexed the reads using the BC tags extracted with the pysam library (http://pysam.readthedocs.io/en/latest/api.html). For bipartite graph display, we used a subset of 2000 BMMC cells and 3300 PBMC cells with the highest number of reads in each dataset, respectively.

### SNV simulation

We created a modified version of the hg19 reference genome by introducing 50000 random mutations in the exonic region of the genes. To introduce a new mutation, we weighted each exon depending on its base length and selected one randomly and proportional to its weight. We realigned the scRNA-seq reads from a subset of 20 cells from the Kim dataset with the introduced new mutations, using the same SNV detection pipelines described earlier. We used BedTools^65^ to compute the read depth of each mutation.

### SNV annotation

To annotate human SNV datasets, dbSNP138 from the NCBI Single Nucleotide Polymorphism database^66^ and reference INDELs from 1000 genomes (1000_phase1 as Mills_and_1000G_gold_standard)^67^ were used. To annotate the mouse SNV dataset, dbSNPv137 for SNPs and INDELs were downloaded from the Mouse Genomes Project of the Sanger Institute, using the following link: ftp://ftp-mouse.sanger.ac.uk/REL-1303-SNPs_Indels-GRCm38/^68^. The mouse SNP databases were sorted using SortVcf command of Picard Tools in order to be properly used by Picard Tools and GATK.

### SSrGE package to correlate eeSNVs to gene expression

For each dataset, we denote *M*_*SNV*_ and *M*_*GE*_ as the SNV and gene expression matrices, respectively. *M*_*SNV*_ is binary 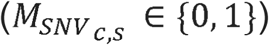 indicating the presence/absence of SNV *s* in cell *c*. 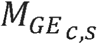 is the log transformed gene expression value of the gene *g* in cell *c*. Copy number variation (CNV) *M*_*CNV*_, can be added as an additional optional predictor in SSrGE. We computed CNV for each gene *g* in each cell *c* using the online platform Ginkgo^69^. For the Hou dataset, we inferred CNV using the same approach described by the authors^43^. We removed any SNV present in less than k=3 cells, as it offers the best tradeoff between accuracy and SSrGE running time. We also filtered SNVs associated with genes having normalized expression value below 2.0. We also discarded genes with normalized expressions below 2.0 and expressed in less than 10 cells from SsrGE analysis. For each gene *g*, we applied a sparse linear regression using LASSO to identify *W*_*g*_, the linear coefficient associated to SNV, as well as W_*cnv*_, the coefficient associated to the CNV of *g* (if CNV was considered). The objective function for SSrGE to minimize is:

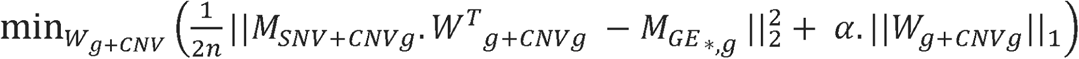

where *α* is the regularization parameter, *M*_*SNV*+*CNV g*_ represents the *M*_*SNV*_ matrix with an additional column that corresponds to the CNV of the gene *g* for each cell. In this configuration, *W*_*g+CNV*_ has one more column too, corresponding to the weight associated with the CNV. An SNV was considered as eeSNV when *W*_*g*_(*s*) ≠ 0. When CNV is not considered in SSrGE, the objective function is simplified as:

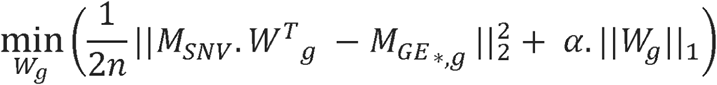

### Inference of SNV genotype and allele-specific SSrGE calibration

We used the package QuASAR (https://github.com/piquelab/QuASAR) to identify the genotype of each SNV^29^. For each dataset, we collected the SNVs and constructed an *n* x 3 matrix using the number of reads mapping to the reference allele, the alternate allele, or neither of the alleles. We then fit this matrix with QuASAR to estimate the true genotype of each SNV. We then estimated the allelic gene expression for each cell by multiplying the normalized gene counts with the fraction of the SNV of a particular genotype. To calibrate SSrGE model with allele expression, we first fit an SSrGE model for each genotype using the allele-specific SNVs and gene expression as inputs. We then merged the eeSNVs and weights inferred for each model into a final model.

### Ranking of eeSNVs and genes

SSrGE generates coefficients of eeSNVs for each gene, as a metric for their contributions to the gene expression. The score of an eeSNV is given by the sum of its weights over all genes:

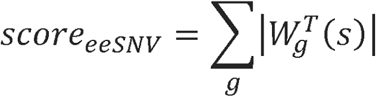

Each gene also receives a score according to its associated eeSNVs:

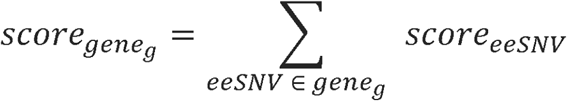

In practice, we first obtained eeSNVs using a minimum filtering of *α*=0.1, before using these two scores above to rank eeSNVs and the genes.

### Ranking of eeSNVs and genes for a subpopulation

For a given single-cell subpopulation *p*, an eeSNV is defined as *specific* to the subpopulation *p* when it has a significantly higher frequency in *p* than in any other subpopulation. For each eeSNV we took only the subset of cells expressing the gene *g* associated with the eeSNV. We then computed the Fisher’s exact test to compare the presence of the eeSNV between single-cells inside and outside *p*. We considered an eeSNV significant for p-value < 0.05. *p’* designates the subset of cells from *p* expressing *g.* The score of an eeSNV for *p* is given by:

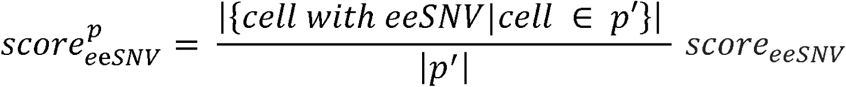

The score of a given gene *g* for *p* is thus given by:

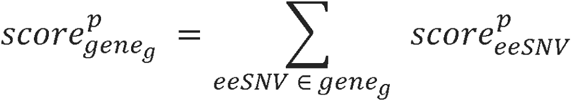

To rank eeSNVs from the promoter regions of the RRBS reads in Hou dataset, we applied a similar methodology: we annotated the eeSNVs within 1500bp upstream of genes’ starting codon regions.

### Perturbation-model based simulation to quantitatively assess SSrGE

We first simulated an interaction table which gives an “interaction score” (between −1 and 1) for each gene-SNV pair, denoted as *interaction* ***gv*** for gene ***g*** and SNV ***v***. This interaction score follows a mixed normal distribution, with two normal distribution components. These two distributions are named the inhibiting component (centered at −0.5), and the enhancing component (centered at 0.5). The type of interaction determines the weighting factor. We define a cis-interaction if the SNV is located within the gene and a trans-interaction otherwise. A cis-interaction has equal weights on the inhibiting and enhancing components. A trans-interaction has significantly larger weight on the inhibiting component, as previous studies found cis interaction is more likely to be inhibitive.

We then simulated the unperturbed expression matrix using Splatter^70^, with parameters estimated from the Kim dataset. We also simulated the SNV matrix using random shuffle of the SNV matrix extracted from the same dataset. We then applied the interactions to the unperturbed expression. For a gene *g* in a sample *s*, the expression level *GE*_*gs*_ was perturbed to *GE’*_*gs*_ using the following formula:

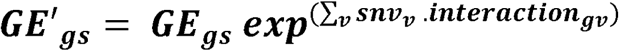

This created the perturbed expression matrix. Then, we added drop-outs to the perturbed expression matrix using different strategies, in order to create the final observed expression matrix. To mimic dropouts, we introduced two variables: one for dropout bias (either bias towards lower expressed genes or randomly), and the other for dropout rate dependency (cell-based, gene-based, or read-based). We did not consider dropout bias towards highly expressed genes, as it contradicts observations documented earlier^71^. Random dropout (no bias) removes the dependency on cell, gene or reads. We also added random noises to the SNV matrix (following a Bernoulli distribution) to generate the final observed SNV matrix.

### Pseudo-time ordering reconstruction

To estimate the trajectory of cell evolvement, we adopted the following procedure, motivated by the method described earlier^38^. We first constructed the following distance matrix to reflect the Pearson’s correlation between each pair of cells: *D*_*i,j*_ *=* 1 *- Correlation*(*Sample*_*i*_, *Sample*_*j*_)

### Subpopulation clustering algorithms

We combined two dimension reduction algorithms: Principal Component Analysis (PCA)^30^ and Factor Analysis (FA)^31^ with two popular clustering approaches: the K-Means algorithm^72^ and agglomerative hierarchical clustering (agglo) with WARD linkage^32^. We also used SIMLR, a recent algorithm specifically tailored to cluster and visualize scRNA-seq data, which learns the similarity matrix from subpopulations^33^. Similar to the original SIMLR study, we used the embedding of the cells produced by the algorithm to apply K-Means algorithm.

PCA and FA were performed using their corresponding implementation in Scikit-Learn (*sklearn*)^73^. For PCA, FA and SIMLR, we used various input dimensions *D* [2, 3, 5, 10, 15, 20, 25, 30] to project the data. To cluster the data with K-Means or the hierarchical agglomerative procedure, we used a different cluster numbers *N* (2 to 80) to obtain the best clustering results from each dataset. We computed accuracy metrics for each (*D N*) pair and chose the combination that gives the overall best score. Between the two clustering methods, K-Means was the implementation of *sklearn* package with the default parameter, and hierarchical clustering was done by the *AgglomerativeClustering* implementation of *sklearn*, using WARD linkage.

### Validation metrics

To assess the accuracy of the obtained clusters, we used three metrics: Adjusted Mutual Information (AMI), Adjusted Rand Index (ARI) and V-measure ^34,35^. These metrics compare the obtained clusters *C* to some reference classes *K* and generate scores between 0 and 1 for AMI and V-measure, and between −1 and 1 for ARI. A score of 1 means perfect match between the obtained clusters and the reference classes. For ARI, a score below 0 indicates a random clustering.

AMI normalizes Mutual Information (MI) against chances^34^. The Mutual Information between two sets of classes C and K is equal to: 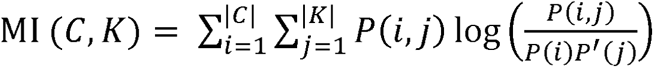, where *p*(*i*) is the probability that an object from *C* belongs to the class *i P′* (*j*) is the probability that an object from *K* belongs to class *j,* and *P*(*i, j*) is the probability that an object are in both class *i* and *j*. AMI is equal to: 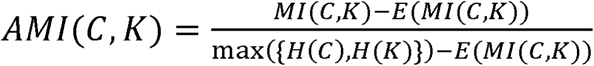, where *H*(*C*) and *H*(*K*) designates the entropy of *C* and *K*.

Similar to AMI, ARI normalizes RI against random chances: 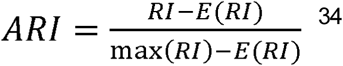 ^34^. Rand Index (RI) was computed by: 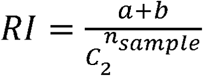, where *a* is the number of con-concordant sample pairs in obtained clusters *C* and reference classes *K*, whereas *b* is the number of dis-concordant samples.

V-measure, similar to F-measure, calculates the harmonic mean between homogeneity and completeness. Homogeneity is defined as 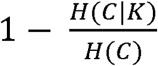, where *H*(*C* | *K*) is the conditional entropy of *C* given *K*. Completeness is the symmetrical of homogeneity: 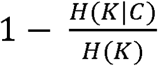.

### Graph visualization

The different datasets were transformed into GraphML files with Python scripts using iGraph library^74^. Graphs were visualized using GePhi software^36^ and spatialized using ForceAtlas2^75^, a specific graph layout implemented into the GePhi software.

### Pathway enrichment analysis

We used the KEGG pathway database to identify pathways related to specific genes^76^. We first selected the top 100 genes for each dataset, according to the ranks given by SSrGE. We then selected genes scored with significant eeSNVs for the metastasis cells from Kim dataset. We used DAVID 6.8 functional annotation tool to identify significant pathways amongst these genes^37^. We used the default significance value (adjusted p-value threshold of 0.10). Significant pathways are represented as a bipartite graph using Gephi: Nodes are either genes or pathway and the size of each nodes represent the score of the genes or, in the case of pathways, the sum of the scores of the genes linked to the pathways. We used the same methodology to infer significant pathways of cancer cells, compared to normal cells, from Hou dataset. However, we used all the genes ranked rather than only the significant genes, since only few genes are found to be significant for cancer cells.

### Code availability

The SNV calling pipeline and SSrGE are available through the following GitHub project: https://github.com/lanagarmire/SSrGE.

## Acknowledgements

This research was supported by grants K01ES025434 awarded by NIEHS through funds provided by the trans-NIH Big Data to Knowledge (BD2K) initiative (www.bd2k.nih.gov), P20 COBRE GM103457 awarded by NIH/NIGMS, R01 LM012373 awarded by NLM, R01 HD084633 awarded by NICHD and Hawaii Community Foundation Medical Research Grant 14ADVC-64566 to L.X. Garmire. We acknowledge K. Chaudhary and John Breck Yunits for manuscript proofreading.

## Author contributions

LG envisioned this project. OP implemented the project and conducted genomics analysis, XZ and TC helped on implementation. OP and LG wrote the manuscript. All authors have read and agreed on the manuscript.

## Competing financial interests

The author(s) declare no competing financial interests.

## Supplemental Materials

**Supplementary Figure S1:** The SNV calling pipeline based on GATK. It follows the “Best Practice” workflow for SNP and INDEL calling as recommended, with four steps. Step 1: alignment. Step 2: preprocessing of BAM files. Step 3: read realignment and recalibration. Step 4: variant calling.

**Supplementary Figure S2:** Performance comparison of GATK vs. FreeBayes SNV calling pipeline using a modified reference h19 genomes containing 50000 random mutations in the exonic region. (A-B) Box plots using a subset of 20 cells from the Kim dataset: (A) true positive rate; (B) false positive rate. (C-D) Box plots using a subset of 100 cells from a 10X genomic dataset: (C) true positive rate; (D) false positive rate.

**Supplementary Figure S3:** Sketch of Sparse SNV inference to Reflect Gene Expression (SSrGE) linear models. The SNVs can be calculated from the GATK pipeline (Supplementary Figure S1) or another SNV calling pipeline preferred by users. These SNVs are transformed into a predictor matrix *M*_*SNV*_. As an option, the users can also include a CNV matrix *M*_*CNV*_ as an additional predictor matrix. CNVs can be inferred from scRNA-Seq data using programs such as the online platform Ginkgo^69^. Gene expression is the response matrix *M*_*GE*_. For each gene, a LASSO regression is fitted to identify non-null coefficient matrix *W*. The output of the models is a set of filtered eeSNVs and a set of corresponding genes in which eeSNVs are found.

**Supplementary Figure S4:** perturbation-model based simulation to evaluate SSrGE quantitatively. Comparisons are performed between the average expected ranks from top genes inferred by SSrGE (x-axis) vs. those set by the simulation (y-axis). (A) Effect of different noise levels. (B) Effect of dropout bias and dropout rate dependency. For dropout bias, it is either biased towards lower expressed genes or random (no bias). Dropout rate is dependent on cell, gene, or reads.

**Supplementary Figure S5**: Relationship between the best accuracy metrics and the LASSO regularization parameter *a*, over the five datasets and five different clustering approaches. The accuracy metrics are: (A) Adjusted Mutual Information (AMI), B: Adjusted Rand Index (ARI), and (C): V-measure.

**Supplementary Figure S6:** Bar plot comparing the clustering performance using eeSNV vs. gene expression (GE) as features, over five datasets and five different clustering strategies. The metrics used are (A): Adjusted Rand Index (ARI), and (B): V-measure.

**Supplementary Figure S7:** Pseudo-time reconstruction using the Monocle algorithm with gene expression features from genes having eeSNVs, as compared to the pseudo-time reconstruction using eeSNVs in the same genes shown in Figure 6A.

**Supplementary Figure S8**: Immune (blue), primary (red) and metastatic (green) tumor cell subpopulations from two breast cancer patients (BC03 and BC07) using either bipartite graphs or minimum spanning trees (Chung dataset).

**Supplementary Figure S9:** Comparison of the batch-effect on SNVs and gene expression, using scRNA-seq data from glioblastoma patient MGH26.

**Supplementary Figure S10**: eeSNV inference, gene ranking, and cell visualization for 9 additional datasets. Datasets E and G are 10X Genomic datasets from the 10X Genomic database website: https://support.10xgenomics.com/single-cell-gene-expression. The other datasets A, B, C, D, F, H, and I are extracted from NCBI GEO. The top 8 genes are presented for each dataset, based on their eeSNV scores.

**Supplementary Figure S11**: Minimum Spanning tree using either eeSNVs (A) or gene expression (B) of HLA as features. Pearson correlation is used as the distance metric. Color labels: primary tumor (green), metastatic cells from patients (blue) and metastatic cells from patient derived xenografts (red).

**Supplementary Table S1:** Genotype of SNVs and similarity testing results with and without allelic specific expression. Average per-cell genotypes of the SNVs are detected and computed using QuASAR. *g0 g1* and *g2* correspond to the *homozygous reference heterozygous,* and *homozygous alternate* genotypes, respectively.

| Dataset | ranked genes<br>($\alpha=0.1$) | | ranked eeSNVs<br>($\alpha=0.1$) | | Number of eeSNVs per genotype | | | |
| --- | --- | --- | --- | --- | --- | --- | --- | --- |
|  | Kendall<br>Tau score | Kendall<br>Tau p-value | Kendall<br>Tau score | Kendall<br>Tau p-value | g0<br>(homo<br>ref) | g1<br>(heter) | g2<br>(homo<br>alter) | Without specific<br>allelic expression |
| Kim | 0.71 | 0 | 0.58 | 0 | 1994 | 4314 | 11876 | 13939 |
| Patel | 0.73 | 0 | 0.62 | 0 | 372 | 2020 | 6660 | 7168 |
| Ting | 0.64 | 0 | 0.53 | 0 | 248 | 2016 | 14214 | 15140 |
| Miyamoto | 0.68 | 0 | 0.58 | 0 | 101 | 1186 | 4981 | 5327 |
| Chung | 0.76 | 0 | 0.53 | 0 | 3026 | 6391 | 16093 | 18741 |

**Supplementary Table S2:** Regularization values (*a*) used for the clustering procedures along with the number of eeSNVs features.

**Supplementary Table S3:** Ranked eeSNVs and genes for each dataset (with minimum regularization filtering *a*=0.1).

**Supplementary Table S4:** Ranked genes for the metastasis single-cells from the Kim dataset (mRCC) and from BC03M and BC07M from Chung datasets.

## Bibliography

1. Harris, K. et al. Molecular organization of CA1 interneuron classes. bioRxiv 34595 (2015).

2. Pierson, E. & Yau, C. ZIFA: Dimensionality reduction for zero-inflated single-cell gene expression analysis. Genome Biol. 16, 1–10 (2015).

3. Vallejos, C. A., Marioni, J. C. & Richardson, S. BASiCS: Bayesian analysis of single-cell sequencing data. PLoS Comput Biol 11, e1004333 (2015).

4. Kolodziejczyk, A. A., Kim, J. K., Svensson, V., Marioni, J. C. & Teichmann, S. A. The technology and biology of single-cell RNA sequencing. Mol. Cell 58, 610–620 (2015).

5. Stegle, O., Teichmann, S. A. & Marioni, J. C. Computational and analytical challenges in single-cell transcriptomics. Nat. Rev. Genet. 16, 133–145 (2015).

6. Buettner, F. et al. Computational analysis of cell-to-cell heterogeneity in single-cell RNA-sequencing data reveals hidden subpopulations of cells. Nat. Biotechnol. 33, 155–160 (2015).

7. Poirion, O. B., Zhu, X., Ching, T. & Garmire, L. Single-cell transcriptomics bioinformatics and computational challenges. Front. Genet. 7, 163 (2016).

8. Bryois, J. et al. Cis and trans effects of human genomic variants on gene expression. PLoS Genet 10, e1004461 (2014).

9. Hu, P., Lan, H., Xu, W., Beyene, J. & Greenwood, C. M. T. Identifying cis-and trans-acting single-nucleotide polymorphisms controlling lymphocyte gene expression in humans. in BMC proceedings 1, 1 (2007).

10. Berdasco, M. & Esteller, M. Aberrant epigenetic landscape in cancer: how cellular identity goes awry. Dev. Cell 19, 698–711 (2010).

11. Navin, N. et al. Tumour evolution inferred by single-cell sequencing. Nature 472, 90–94 (2011).

12. Almendro, V., Marusyk, A. & Polyak, K. Cellular heterogeneity and molecular evolution in cancer. Annu. Rev. Pathol. Mech. Dis. 8, 277–302 (2013).

13. Burrell, R. A., McGranahan, N., Bartek, J. & Swanton, C. The causes and consequences of genetic heterogeneity in cancer evolution. Nature 501, 338–345 (2013).

14. Zafar, H., Wang, Y., Nakhleh, L., Navin, N. & Chen, K. Monovar: single-nucleotide variant detection in single cells. Nat. Methods (2016).

15. Ross, E. M. & Markowetz, F. OncoNEM: inferring tumor evolution from single-cell sequencing data. Genome Biol. 17, 1 (2016).

16. Welch, J. D., Hartemink, A. J. & Prins, J. F. SLICER: inferring branched, nonlinear cellular trajectories from single cell RNA-seq data. Genome Biol. 17, 1 (2016).

17. Gamazon, E. R. et al. PrediXcan: Trait Mapping Using Human Transcriptome Regulation. bioRxiv 20164 (2015).

18. Pineda, S. et al. Integration analysis of three omics data using penalized regression methods: An application to bladder cancer. PLoS Genet 11, e1005689 (2015).

19. Ortega, M. A. et al. Using single-cell multiple omics approaches to resolve tumor heterogeneity. Clin. Transl. Med. 6, 46 (2017).

20. Kim, K.-T. et al. Application of single-cell RNA sequencing in optimizing a combinatorial therapeutic strategy in metastatic renal cell carcinoma. Genome Biol. 17, 80 (2016).

21. Ting, D. T. et al. Single-cell RNA sequencing identifies extracellular matrix gene expression by pancreatic circulating tumor cells. Cell Rep. 8, 1905–1918 (2014).

22. Miyamoto, D. T. et al. RNA-Seq of single prostate CTCs implicates noncanonical Wnt signaling in antiandrogen resistance. Science (80-.). 349, 1351–1356 (2015).

23. Patel, A. P. et al. Single-cell RNA-seq highlights intratumoral heterogeneity in primary glioblastoma. Science (80-.). 344, 1396–1401 (2014).

24. Chung, W. et al. Single-cell RNA-seq enables comprehensive tumour and immune cell profiling in primary breast cancer. Nat. Commun. 8, (2017).

25. Piskol, R., Ramaswami, G. & Li, J. B. Reliable identification of genomic variants from RNA-seq data. Am. J. Hum. Genet. 93, 641–651 (2013).

26. Garrison, E. & Marth, G. Haplotype-based variant detection from short-read sequencing. arXiv Prepr. arXiv1207.3907 (2012).

27. Tibshirani, R. Regression shrinkage and selection via the lasso. J. R. Stat. Soc. Ser. B 267–288 (1996).

28. Nelsen, R. B. Kendall tau metric. Encycl. Math. 3, 226–227 (2001).

29. Harvey, C. T. et al. QuASAR: quantitative allele-specific analysis of reads. Bioinformatics 31, 1235–1242 (2014).

30. I T, J. ‘Principal Component Analysis, 2nd ed’. Journal of the American Statistical Association 98, (Springer Series in Statistics, 2002).

31. Cattell, R. B. Factor analysis: an introduction and manual for the psychologist and social scientist. (1952).

32. Joe H Ward, J. Hierarchical Grouping to Optimize an Objective Function. J. Am. Stat. Assoc. 48, 236–244 (1963).

33. Wang, B., Zhu, J., Pierson, E. & Batzoglou, S. Visualization and analysis of single-cell RNA-seq data by kernel-based similarity learning. bioRxiv 52225 (2016).

34. Vinh, N. X., Epps, J. & Bailey, J. Information theoretic measures for clusterings comparison: Variants, properties, normalization and correction for chance. J. Mach. Learn. Res. 11, 2837–2854 (2010).

35. Rosenberg, A. & Hirschberg, J. V-Measure: A conditional entropy-based external cluster evaluation measure. Comput. Linguist. 410–420 (2007).

36. Bastian, M., Heymann, S. & Jacomy, M. Gephi: An open source software for exploring and manipulating networks. (2009).

37. Huang, D. W., Sherman, B. T. & Lempicki, R. A. Systematic and integrative analysis of large gene lists using DAVID bioinformatics resources. Nat. Protoc. 4, 44–57 (2009).

38. Trapnell, C. et al. Pseudo-temporal ordering of individual cells reveals dynamics and regulators of cell fate decisions. Nat. Biotechnol. 32, 381 (2014).

39. Kuriyama, S. et al. LPP inhibits collective cell migration during lung cancer dissemination. Oncogene 35, 952–964 (2016).

40. Fedele, M. et al. Role of the high mobility group A proteins in human lipomas. Carcinogenesis 22, 1583–1591 (2001).

41. Godar, S. et al. Growth-inhibitory and tumor-suppressive functions of p53 depend on its repression of CD44 expression. Cell 134, 62–73 (2008).

42. Wielenga, V. J. M. et al. Expression of CD44 variant proteins in human colorectal cancer is related to tumor progression. Cancer Res. 53, 4754–4756 (1993).

43. Hou, Y. et al. Single-cell triple omics sequencing reveals genetic, epigenetic, and transcriptomic heterogeneity in hepatocellular carcinomas. Cell Res. 26, 304–319 (2016).

44. Oh, T. G. et al. PRMT2 and ROR$*γ*$ expression are associated with breast cancer survival outcomes. Mol. Endocrinol. 28, 1166–1185 (2014).

45. Lai, J.-P. et al. Sulfatase 2 protects hepatocellular carcinoma cells against apoptosis induced by the PI3K inhibitor LY294002 and ERK and JNK kinase inhibitors. Liver Int. 30, 1522–1528 (2010).

46. Tirosh, I. et al. Dissecting the multicellular ecosystem of metastatic melanoma by single-cell RNA-seq. Science (80-.). 352, 189–196 (2016).

47. de Bakker, P. I. W. et al. A high-resolution HLA and SNP haplotype map for disease association studies in the extended human MHC. Nat. Genet. 38, 1166–1172 (2006).

48. Network, C. G. A. R. & others. Comprehensive molecular characterization of gastric adenocarcinoma. Nature 513, 202–209 (2014).

49. Shukla, S. A. et al. Comprehensive analysis of cancer-associated somatic mutations in class I HLA genes. Nat. Biotechnol. 33, 1152–1158 (2015).

50. Chang, C.-C., Campoli, M., Restifo, N. P., Wang, X. & Ferrone, S. Immune selection of hot-spot $ß$2-microglobulin gene mutations, HLA-A2 allospecificity loss, and antigen-processing machinery component down-regulation in melanoma cells derived from recurrent metastases following immunotherapy. J. Immunol. 174, 1462–1471 (2005).

51. Sveen, A., Kilpinen, S., Ruusulehto, A., Lothe, R. A. & Skotheim, R. I. Aberrant RNA splicing in cancer; expression changes and driver mutations of splicing factor genes. Oncogene 35, 2413–2428 (2016).

52. Macaulay, I. C. et al. G&T-seq: parallel sequencing of single-cell genomes and transcriptomes. Nat. Methods 12, 519–522 (2015).

53. Shannon, P. et al. Cytoscape: a software environment for integrated models of biomolecular interaction networks. Genome Res. 13, 2498–2504 (2003).

54. Dey, S. S., Kester, L., Spanjaard, B., Bienko, M. & van Oudenaarden, A. Integrated genome and transcriptome sequencing of the same cell. Nat. Biotechnol. 33, 285–289 (2015).

55. Kim, K.-T. et al. Single-cell mRNA sequencing identifies subclonal heterogeneity in anti-cancer drug responses of lung adenocarcinoma cells. Genome Biol 16, 127 (2015).

56. Zhu, X. et al. Granatum: a graphical single-cell RNA-Seq analysis pipeline for genomics scientists. Genome Med. 9, 108 (2017).

57. Barrett, T. et al. NCBI GEO: archive for functional genomics data sets—update. Nucleic Acids Res. 41, D991--D995 (2013).

58. Auwera, G. A. et al. From FastQ data to high-confidence variant calls: the genome analysis toolkit best practices pipeline. Curr. Protoc. Bioinforma. 10–11 (2013).

59. Dobin, A. & Gingeras T. R. Mapping RNA-seq Reads with STAR. Curr. Protoc. Bioinforma. 11–14 (2015).

60. Andrews, S. & others. FastQC: A quality control tool for high throughput sequence data. Ref. Source (2010).

61. Liao, Y., Smyth, G. K. & Shi, W. featureCounts: an efficient general purpose program for assigning sequence reads to genomic features. Bioinformatics btt656 (2013).

62. Guidot, A. et al. Genomic structure and phylogeny of the plant pathogen Ralstonia solanacearum inferred from gene distribution analysis. J. Bacteriol. 189, 377–87 (2007).

63. Krueger, F. & Andrews, S. R. Bismark: a flexible aligner and methylation caller for Bisulfite-Seq applications. bioinformatics 27, 1571–1572 (2011).

64. Gao, S. et al. BS-SNPer: SNP calling in bisulfite-seq data. Bioinformatics btv507 (2015).

65. Quinlan, A. R. & Hall, I. M. BEDTools: a flexible suite of utilities for comparing genomic features. Bioinformatics 26, 841–842 (2010).

66. Sherry, S. T. et al. dbSNP: the NCBI database of genetic variation. Nucleic Acids Res. 29, 308–311 (2001).

67. Consortium, 1000 Genomes Project & others. A global reference for human genetic variation. Nature 526, 68–74 (2015).

68. Keane, T. M. et al. Mouse genomic variation and its effect on phenotypes and gene regulation. Nature 477, 289–294 (2011).

69. Garvin, T. et al. Interactive analysis and assessment of single-cell copy-number variations. Nat. Methods 12, 1058 (2015).

70. Zappia, L., Phipson, B. & Oshlack, A. Splatter: Simulation Of Single-Cell RNA Sequencing Data. bioRxiv 133173(2017).

71. Brennecke, P. et al. Accounting for technical noise in single-cell RNA-seq experiments. Nat. Methods 10, 1093–1095 (2013).

72. MacQueen, J. & others. Some methods for classification and analysis of multivariate observations. in Proceedings of the fifth Berkeley symposium on mathematical statistics and probability 1, 281–297 (1967).

73. Pedregosa, F., Weiss, R. & Brucher, M. Scikit-learn[: Machine Learning in Python. J. Mach. Learn. Res. 12, 2825–2830 (2011).

74. Csardi, G. & Nepusz, T. The igraph software package for complex network research. InterJournal Complex Syst. Complex Sy, 1695 (2006).

75. Jacomy, M., Venturini, T. & Bastian, M. ForceAtlas2, A Graph Layout Algorithm for Handy Network Visualization. 1–21 (2011).

76. Kanehisa, M., Sato, Y., Kawashima, M., Furumichi, M. & Tanabe, M. KEGG as a reference resource for gene and protein annotation. Nucleic Acids Res. gkv1070 (2015).

